# Structural Basis for Dimerization and Activation of UvrD-family Helicases

**DOI:** 10.1101/2024.09.05.611425

**Authors:** Ankita Chadda, Binh Nguyen, Timothy M. Lohman, Eric A. Galburt

## Abstract

UvrD-family helicases are superfamily 1A motor proteins that function during DNA replication, recombination, repair, and transcription. UvrD family monomers translocate along single stranded (ss) DNA but need to be activated by dimerization to unwind DNA in the absence of force or accessory factors. However, prior structural studies have only revealed monomeric complexes. Here, we report the first structures of a dimeric UvrD-family helicase, *Mycobacterium tuberculosis* UvrD1, both free and bound to a DNA junction. In each structure, the dimer interface occurs between the 2B subdomains of each subunit. The apo UvrD1 dimer is observed in symmetric compact and extended forms indicating substantial flexibility. This symmetry is broken in the DNA-bound dimer complex with leading and trailing subunits adopting distinct conformations. Biochemical experiments reveal that the *E. coli* UvrD dimer shares the same 2B-2B interface. In contrast to the dimeric structures, an inactive, auto-inhibited UvrD1 DNA-bound monomer structure reveals 2B subdomain-DNA contacts that are likely inhibitory. The major re-orientation of the 2B subdomains that occurs upon UvrD1 dimerization prevents these duplex DNA interactions, thus relieving the auto-inhibition. These structures reveal that the 2B subdomain serves a major regulatory role rather than participating directly in DNA unwinding.

## Introduction

Helicases are ubiquitous enzymes involved in genome maintenance including replication, transcription, translation, recombination, and DNA repair (*1*). These motor proteins use ATP binding and hydrolysis to perform the physical work of DNA translocation and separation (*i*.*e*., unwinding) of the complementary DNA strands (*2-4*). Helicases are divided into six superfamilies (SF). SF3-6 enzymes form hexameric rings and include replicative DNA helicases (*5, 6*). SF1-2 enzymes are involved in DNA repair, replication restart, transcription, chromatin remodeling and RNA maintenance (*7-11*). SF1 enzymes can be sub-divided based on whether they translocate 3’ to 5’ (SF1A) or 5’ to 3’ (SF1B) and the SF1A group contains the UvrD-family, based on homology to the *E. coli* (*Ec*) UvrD enzyme (*7*). UvrD-family helicases participate in a variety of biochemical pathways, including nucleotide excision repair (*12, 13*), DNA mismatch repair (*14*) and transcription-coupled repair (*15*).

Structures of monomeric UvrD-family enzymes have been reported in apo (*16, 17*) and DNA-bound forms (*17-20*). The conserved four subdomain structure of UvrD-family enzymes consists of the RecA-like 1A and 2A motor subdomains, which form an ATP hydrolysis active site at their interface, interrupted by large insertions that form the 1B and 2B subdomains. Previous structures of UvrD-family helicases (Rep, PcrA, and UvrD) show that the 2B subdomain can undergo large rotations about a hinge region connecting it to the 2A subdomain (*16-20*). The 2B subdomain is in a closed conformation in the *Ec*UvrD-DNA structure (*20*) and rotates 160^°^ to an open conformation in the apo *Ec*UvrD structure (*16*). Based on the existence of monomeric *Ec*UvrD-DNA (*20*) and *B. subtilis* PcrA-DNA complex structures (*19*) it has been proposed that the functional UvrD-family helicase is monomeric and that the 2B subdomain plays a direct role in DNA unwinding. However, *Ec*UvrD exists as monomers, dimers, and tetramers (*21*) and biochemical experiments show that the monomeric form of these enzymes, while able to use ATP to translocate directionally along ssDNA (*22-24*), is not able to processively unwind DNA in the absence of force or auxiliary factors (*23, 25-27*). Dimerization of these enzymes is needed for processive DNA unwinding (*26-33*). Furthermore, deletion of the 2B subdomain in *Ec*Rep activates the monomeric helicase suggesting that the 2B subdomain plays an auto-inhibitory role in the monomer rather than a catalytic one (*28, 34, 35*). The SF1A *Mycobacterium tuberculosis* (*Mt*) UvrD1 helicase, which shares 42% sequence identity with *Ec*UvrD, is activated via redox-dependent dimerization (*32*). Here, we report structures of the dimeric form of the *Mt*UvrD1 helicase that suggest a general mechanism for activation and regulation of UvrD-family helicases and translocases.

## Results

### Apo UvrD1 dimers exist in at least two conformations

*Mt*UvrD1 possesses the four canonical subdomains of SF1 enzymes (Fig. 1A) and a disordered C-terminal linker that leads to a Tudor domain (T). We have previously shown that UvrD1 forms a redox-dependent dimer via disulfide bond formation between the same two cysteines (C451) in each 2B subdomain. Furthermore, this dimer, and not the monomer, is a processive helicase (*32*). To investigate the structural mechanism of UvrD1 dimerization, we first obtained structures of dimers in oxidative conditions in the absence of DNA using cryo-electron microscopy single particle reconstruction (Supplemental Methods). 2D classification (Fig. 1B, fig. S1) and 3D particle reconstruction (Fig. 1C, fig. S1) revealed two distinct conformations which we refer to as “compact” and “extended”. Initial alignments with previously determined monomeric structures revealed the general regions of density pertaining to each conserved subdomain (Fig. 1C,D). The compact structure (4 Å overall, fig. S2) exhibits multiple contacts between the two subunits and a 2B-2B interface is observed consistent with the essentiality of the 2B-2B disulfide bond for dimerization (Fig. 1C). In addition, dimer contacts are observed between the two 1A subdomains which results in the compact nature of this conformation. Each of these contacts are described in more detail below. In contrast, the considerably lower resolution structure of the extended dimer is held together exclusively by the 2B-2B interface (5 Å overall, Fig. 1D, fig. S3). The lower resolution and the asymmetry in resolution between subunits suggests that there exists substantial flexibility across the dimer interface and that the two subunits may take on a distribution of conformations (fig. S3 and S4).

**Fig. 1:**
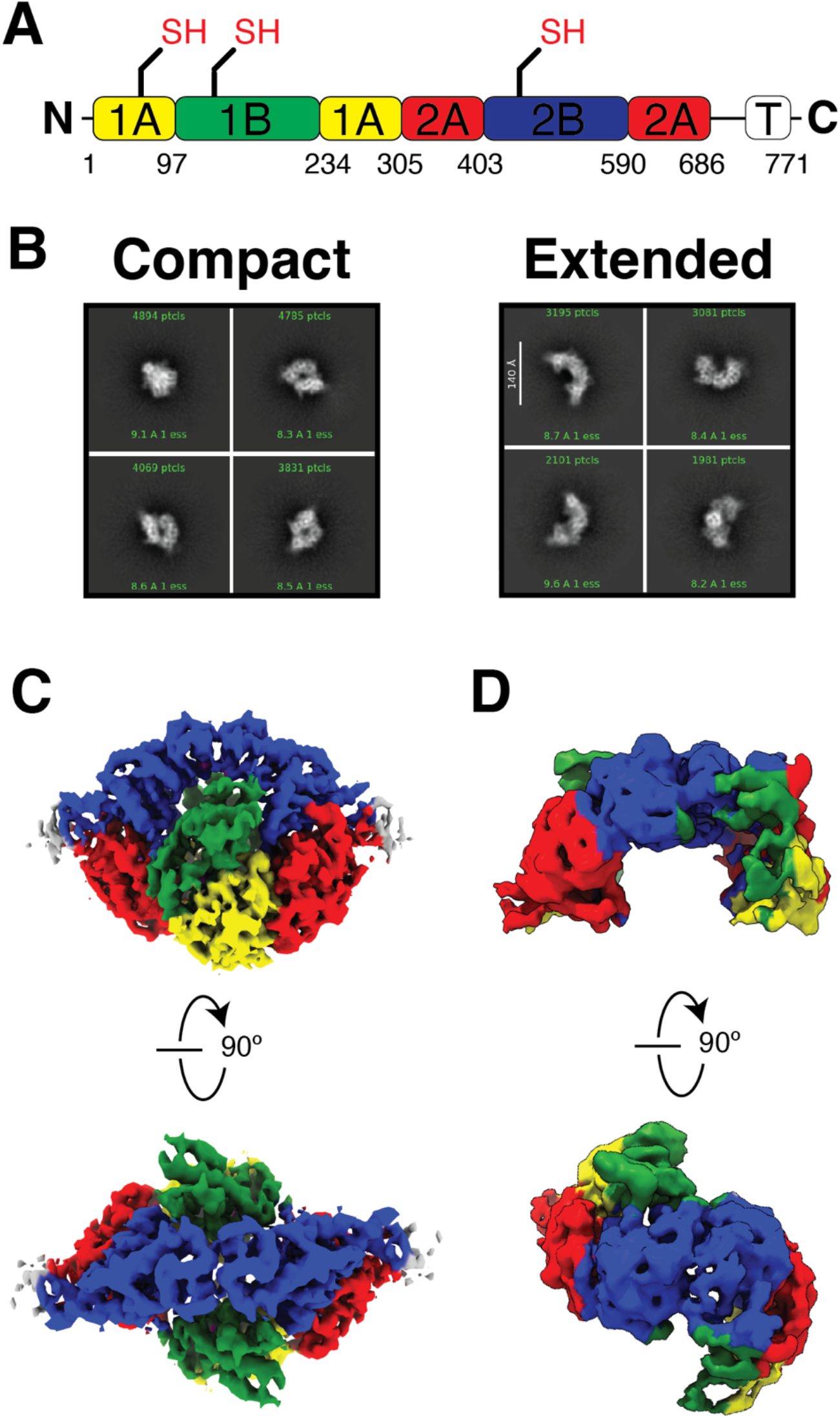
Two conformations of the apo UvrD1 dimer. **(A)** Primary structure and domain organization. **(B)** Selected 2D class averages for compact and extended conformations. **(C and D)** Two views of the compact and extended structure respectively color coded according to the domain colors in (A).

### 2B domain conformation and the 2B-2B dimerization interface

Both the compact and extended conformations of dimeric UvrD1 show the 2B subdomain of each subunit in an open conformation. This can clearly be seen in a structural alignment between one of the subunits of the dimer and the previously determined “open” *Ec*UvrD structure (*16*) fig. S5). The compact structure allows a more precise description of the 2B-2B domain interface (movie S1). In Fig. 2, the cysteines (C451) responsible for redox-dependent dimerization occupy positions consistent with the presence of a disulfide bond. In addition, the sequence C-terminal to C451 (V452-T459, light orange) appears in a strained helical conformation that faces the other subunit and may also contribute to the dimer interface. Continuing towards the C-terminus, density bridging the loops formed by residues K474-N479 (dark orange) suggests additional stabilizing contacts.

**Fig. 2:**
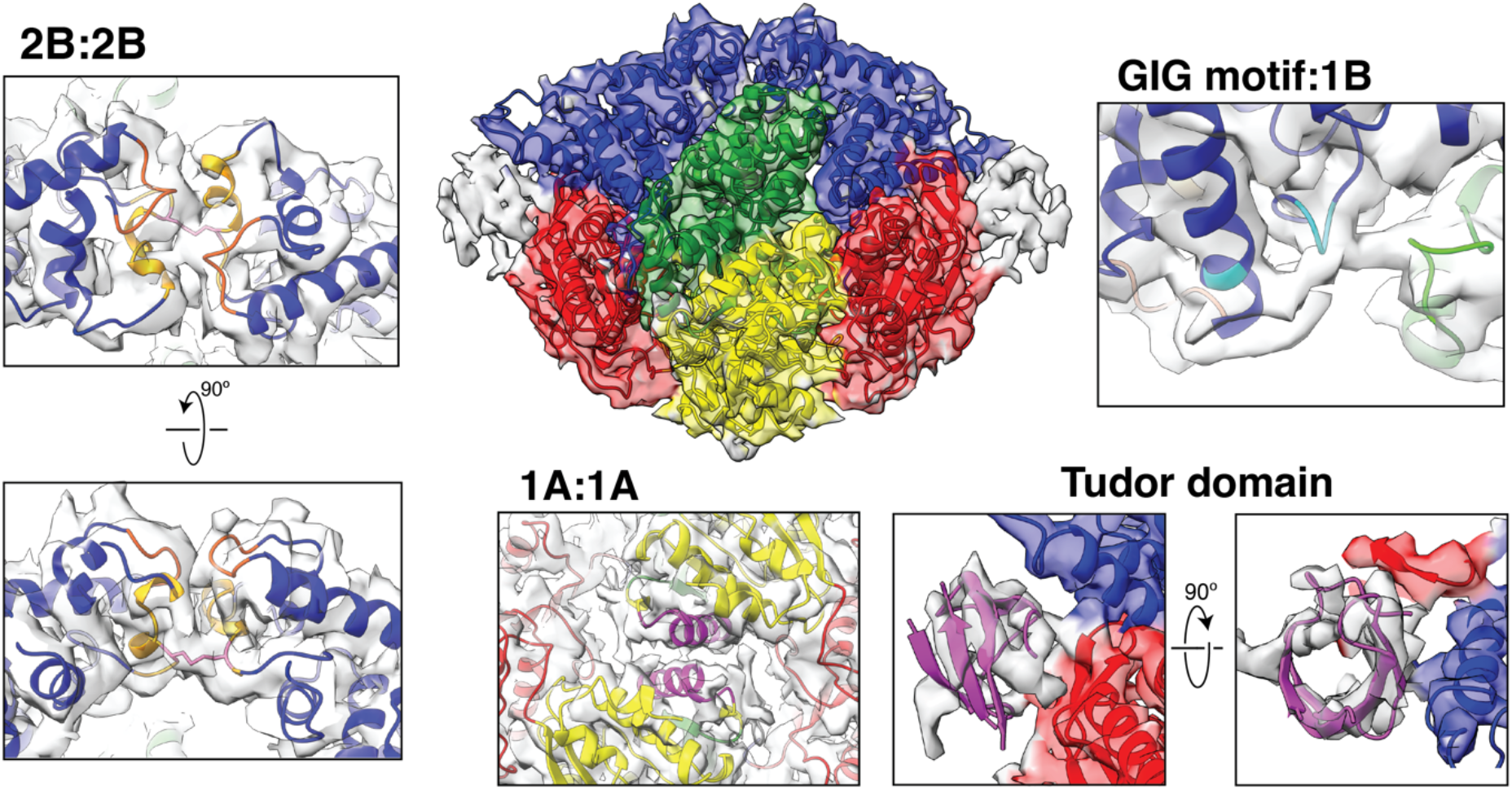
The compact apo structure delineates the 2B-2B dimer interface, other subunit interactions, and a view of the C-terminal Tudor domain. **(2B-2B)** The disulfide bond (pink), the strained helix C-terminal to C451 (light orange), and the loop containing M477 (dark orange) form the interface. **(GIG motif-1B)** The 2B domain (blue) G433-I434-G435 loop and K484 are shown in cyan while the 1B domain loop (L146-R150) is highlighted in green. **(1A-1A)** The 1A domain (yellow) helices (N77-V91) are highlighted in pink. **(Tudor)** AlphaFold model of the UvrD1 Tudor domain (purple) fits the unaccounted-for experimental density nestled between the 2A and 2B subdomains of each subunit.

### Additional contacts in the compact dimer include cis-1B-2B and trans-1A-1A interactions

The compact apo conformation reveals 2 additional contacts of interest. First, within a subunit, the 1B domain contacts the 2B domain in an interaction involving the conserved GIG motif (G443-G445) previously observed contacting duplex DNA in the context of an *Ec*UvrD monomer bound to a ss/ds DNA junction (*20*). Visible links in the density suggests the involvement of 1B K149, the GIG residues, and 2B K484 (Fig. 2, box GIG-motif-1B). Second, 1A helices (N77-V91) come in close proximity to one another across the axis of symmetry with multiple bridging density features (Fig. 2, box 1A-1A).

### The C-terminal Tudor domain

The C-terminal Tudor domain is a beta-stranded structure involved in protein-protein interactions. In the case of UvrD-family enzymes including *Ec*UvrD and *B. subtilis* PcrA, this domain facilitates interactions with RNA polymerase (*36-38*). In the compact UvrD1 conformation, we observe unaccounted for density at the distal ends of the structure at the interface between the 2A and 2B subdomains (Fig. 2). An AlphaFold predicted structure of the UvrD1 Tudor domain fits the size and shape of the density well (Fig. 2, box Tudor). The 38 residues linking the 2A and Tudor domains are not observed consistent with its assignment as a disordered linker. Deletion of the C-terminal 73 amino acids of *Ec*UvrD is known to destabilize UvrD dimers (*27*), which might be explained by domain swapping of the Tudor domains with opposite subunits of the dimer.

### The structure of DNA bound UvrD1 dimer

UvrD1 (6 μM monomer units) under oxidative conditions was incubated with 3’-dT_20_-18bp DNA (6 μM) prior to vitrification and image collection. 2D classification indicated the existence of both apo UvrD1 in a compact conformation along with DNA-bound UvrD1 (fig. S6). Single particle reconstruction of the DNA bound particles resulted in a structure of the dimeric enzyme bound to the ss-dsDNA junction (5.9 Å overall, Fig. 3A, figs. S7-8, movie S2). Although the 2B-2B dimer interface is maintained, an initial survey of the structure reveals a massive conformational change compared to the apo compact structure. In particular, the C_2_ symmetry of the apo dimer has broken, and the conformation of each subunit is distinct. Neither 2B domain is in proximity of the duplex DNA, but a leading and trailing subunit can be identified based on the position of density corresponding to the DNA duplex. The duplex DNA density is of lower local resolution than the rest of the structure (fig. S8) suggesting that it adopts multiple orientations relative to the enzyme. Collecting a larger dataset and masking of the dsDNA region of the map led to a higher resolution map (Supplemental Methods, 4.9 Å overall, figs. 9-10) that more clearly delineates the features of the DNA-bound UvrD1 dimer and the path of the ssDNA (Fig. 3B). For example, both subunits interact with the ssDNA, consistent with previous work showing the requirement of longer ssDNA tails to recruit active dimeric enzymes (*27*).

**Figure 3:**
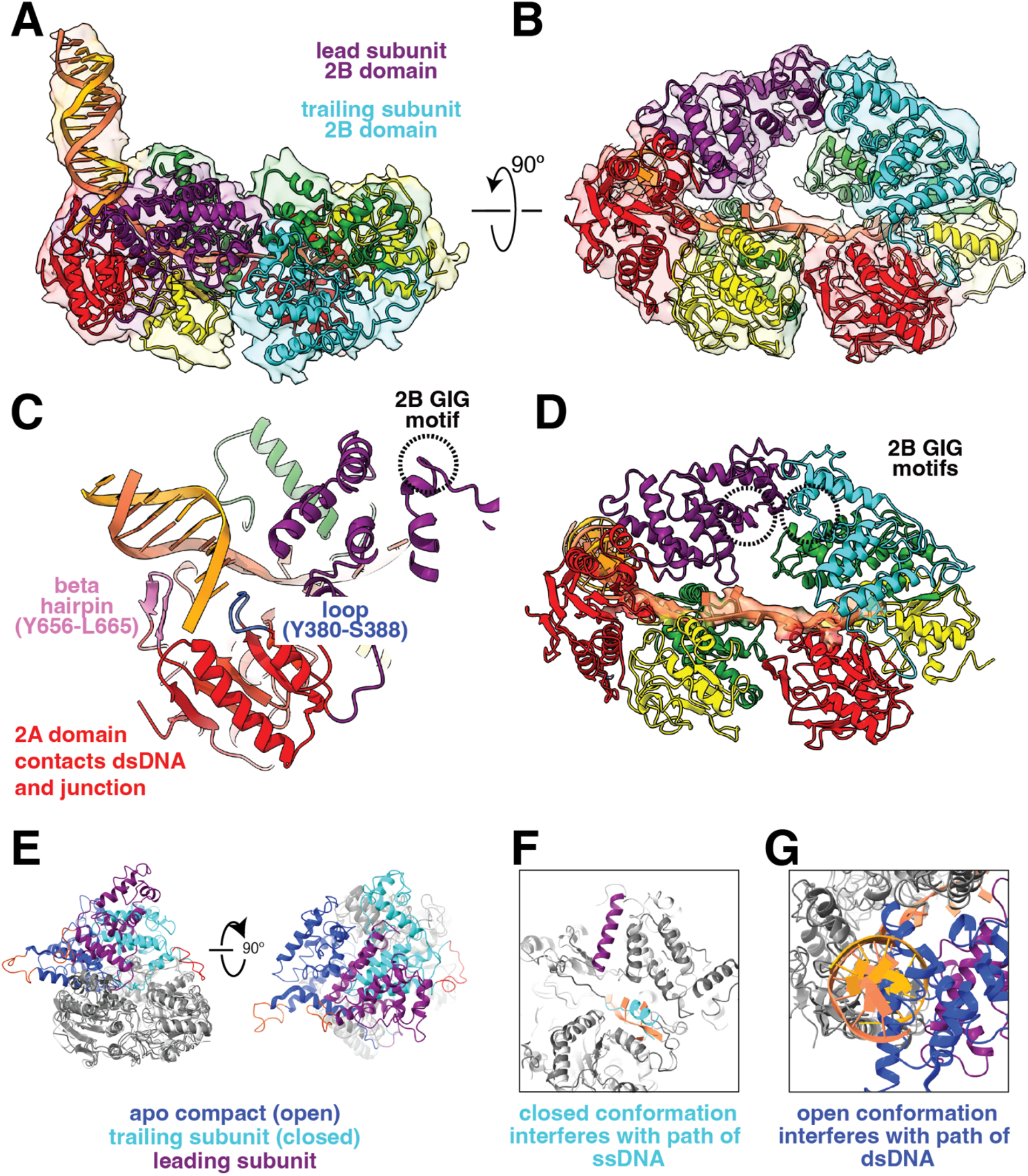
The DNA-bound UvrD1 dimer. **(A)** Unmasked map showing the position of the dsDNA. A leading and trailing subunit were identified based on their position relative to the junction and marked with purple and cyan 2B subdomains respectively. **(B)** Masking the dsDNA resulted in a map that more clearly resolves the protein conformation and the path of the ssDNA. **(C)** The dsDNA is bound primarily by the 2A subdomain of the leading subunit while the 2B GIG motif is far away. A beta hairpin (magenta) and a loop structure (blue) are highlighted. **(D)** The ssDNA interacts with both subunits in a 2A-1A-2A-1A orientation (3’-5’). The 2B GIG motifs are close to the dimer interface and do not contact the DNA. The leading motif (purple) faces the trailing 1B domain and the trailing motif (cyan) faces the solvent. **(E)** Alignment of UvrD1 subunits using the 1A, 1B, and 2A subdomains (greyscale) reveals a variety of 2B subdomain conformations. A loop formed by residues 551-564 (orange) is highlighted as a structural marker of conformation. The leading subunit displays a new 2B domain conformation (purple) which is distinct from that of the trailing subunit (closed, cyan) or the apo compact subunits (open, blue). **(F)** The closed conformation would result in a clash between the 2B “gating helix” (cyan) and the path of the ssDNA through the lead subunit compared to the leading 2B conformation (gating helix in purple). **(G)** The open conformation (blue) would result in a clash with the location of the dsDNA on the lead subunit compared to the leading 2B conformation (purple).

Starting from the DNA junction, there are multiple interactions with the 2A motor subdomain of the leading subunit (Fig. 3C). The beta-hairpin homologous to a region in *Ec*UvrD previously identified as a strand separation element (V656-L665), is found in the major groove of the duplex and instead the loop formed by residues Y380-S388 is positioned where strand separation would be expected to occur (Fig. 3C, fig. S11, and movie S3). Moving towards the 3’ end, the ssDNA passes through the 1A domain while also coming in close proximity with the 1B subdomain (N122-N124, A202-D206 and H104-N133). In the leading subunit, the GIG motif previously observed to interact with duplex DNA in the monomeric *Ec*UvrD and *Bs*PcrA structures (*19, 20*), instead faces a loop in the 1B subdomain of the trailing subunit (N143-L146) (Fig. 3C,D). The ssDNA continues onto the 2A and 1A subdomains of the trailing subunit. Thus, the motor subdomains of each subunit are bound in the same direction with respect to the nucleic acid polarity. This is in contrast to the apo compact structure where the C_2_ symmetry would require a DNA loop for the motor domains to be oriented the same on ssDNA (fig. S12). In the trailing subunit, the GIG motif faces the solvent (Fig. 3D).

While the trailing subunit has a closed 2B conformation (fig. S13), the leading subunit exhibits a previously unobserved conformation (Fig. 3E). Movements of the 2B subdomain are facilitated by hinge regions that link it to the 2A domain and can be quantitated by degrees of rotation around a defined axis. For example, the apo compact (open) and trailing (closed) conformations are related by a rotation of ∼154°. The leading and trailing subunit 2B domains are related by a 147° rotation, but with a distinct axis of rotation (fig. S14, movie S4). The ability of the 2B domain to rotate around different axes broadens the accessible conformational space. Thus, the joint connecting the 2B domain to the rest of the protein may be more analogous to a universal joint than a hinge. Alignment of the trailing subunit with the leading subunit and the junction DNA reveals that a closed conformation would create a clash between the previously identified “gating helix” and the ssDNA as it leaves the leading subunit and transitions towards the trailing subunit (Fig. 3F, movie S5). In addition, alignment of the open apo compact conformation reveals that this conformation would interfere with the position of the duplex DNA held by the leading subunit (Fig. 3G, movie S6). Therefore, rotation of the 2B domain around a unique axis results in a conformation that allows both the 2A and 1A motor domain to interact with duplex DNA and ssDNA, respectively in a manner productive for unwinding.

### Structure of the inhibited *Mt*UvrD1 monomer bound to a DNA junction

With the objective of understanding why monomeric UvrD1 is inhibited for DNA unwinding, we solved a UvrD1 monomer-DNA junction structure using a previously studied constitutively monomeric C451A mutant bound to a 3’-dT_10_-18bp DNA (Supplemental Methods, 5.6 Å overall, figs. S15-16). The density is well accounted for using the AlphaFold prediction for a monomer of UvrD1 aligned with the previously observed closed conformation of a monomer of *Ec*UvrD bound to junction DNA (*20*) (Fig. 4A,B, movie S7). In this structure, the 2B domain (including the GIG motif) are in position to contact the dsDNA as previously observed (*20*). A comparison with the dimeric structure described above indicates that UvrD1 dimerization prevents this interaction as the 2B subdomain is sequestered in the protein-protein interface of the dimer (Fig. 4C, movie S8). Hence, the 2B-DNA interaction and UvrD1 dimerization are competitive and mutually exclusive processes. The 2B subdomain undergoes a large rotation in transitioning from the monomer-DNA complex to the lead subunit-DNA complex in the dimer (Figure 4C, movie S9). In addition, since the DNA-bound monomer conformation is closed (as is the trailing subunit of the DNA-bound dimer), 2B residues 566-580 would also interfere with the path of the ssDNA observed through the leading subunit of the dimeric DNA-bound enzyme (Fig. 3F). Since no helicase activity has been observed for any UvrD-family monomers in the absence of force or accessory proteins, the monomeric 2B subdomain position, its interaction with duplex DNA, and its potential interference with the path of ssDNA are likely inhibitory to helicase activity. Dimerization via the 2B-2B interaction is one mechanism for removing this inhibition.

**Figure 4:**
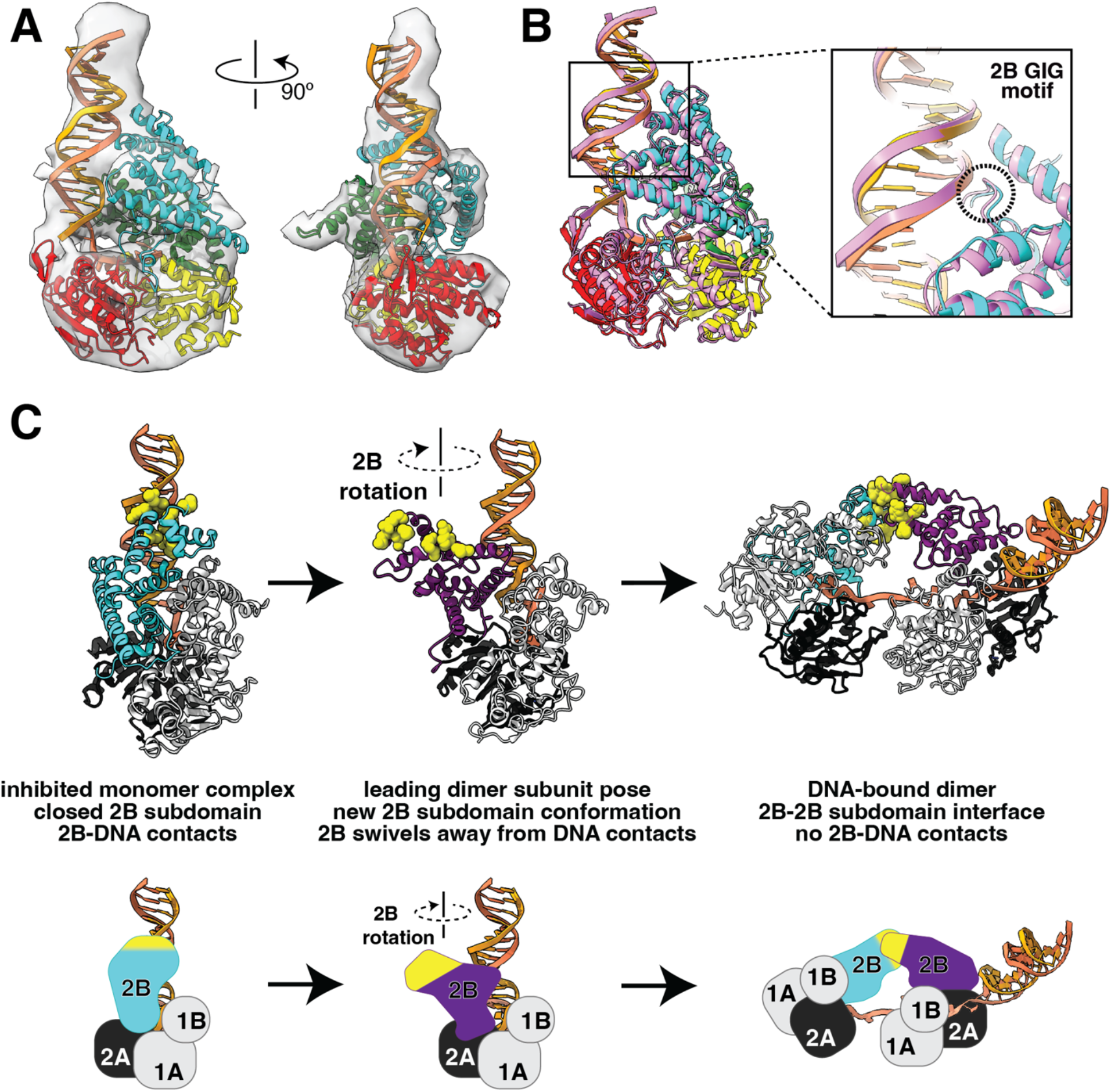
The inactive monomer-DNA complex: **(A)** Experimental density and model for monomeric UvrD1 C451A bound to junction DNA. The 2B domain (cyan) is in the closed conformation. **(B)** Alignment of UvrD1 monomer-DNA complex with E. coli UvrD structure (2IS6, magenta). Inset shows the position of the GIG motif of the 2B domain in each case is in proximity of the dsDNA. **(C)** The closed position of the 2B subdomain in the DNA-bound monomer (cyan) results in contact between the subdomain and duplex DNA (yellow). The 2B subdomain swivels out dramatically from these DNA-contacts in the leading subunit of the dimer and the residues involved now participate in the 2B-2B dimer interface.

### The 2B-2B dimer interface is conserved in *Ec*UvrD

To test whether the UvrD1 dimer interface is conserved in non-disulfide bonded UvrD-family members, we constructed a homology model of an *Ec*UvrD dimer based on the dimeric apo compact UvrD1 structure (Fig. 2 and 5A). This alignment indicates that arginine (R421) in *Ec*UvrD occupies the position of the *Mt*UvrD1 cysteine (C451). If both enzymes use the same dimer interface, mutating the arginine to a cysteine (R421C) should result in a disulfide bond between the two 2B subdomains in *Ec*UvrD under oxidizing conditions. We incorporated this mutation into a Cys-less *Ec*UvrD and used sedimentation velocity experiments to quantify dimerization under both oxidizing and reducing conditions. Consistent with our expectations, at 15 nM total protein in the presence of DTT, *Ec*UvrD(R421C) exists only as a monomer, but in the absence of DTT, ∼41% of the protein forms dimers (Fig. 5B, top panel).

**Figure 5:**
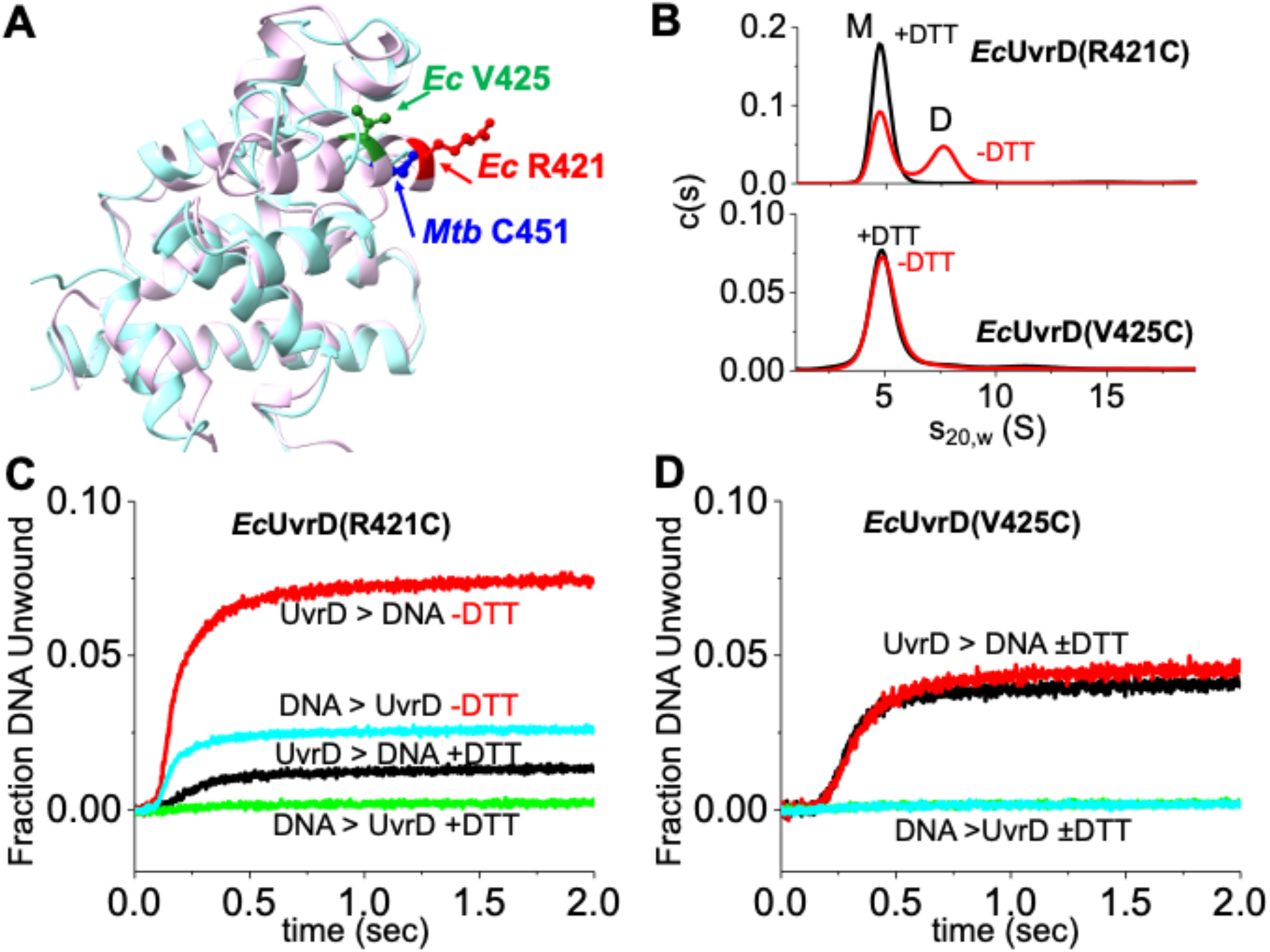
EcUvrD(R421C) forms a redox-dependent dimer which is an active helicase. **(A)** Structural alignment of the 2B subdomains of MtbUvrD1 (cyan) and EcUvrD (purple) shows that MtC451 is structurally equivalent to EcR421 (red). EcV425 (green) is indicated. **(B)** Top: Sedimentation coefficient distribution (c(s)) for 175 nM EcUvrD(R421C) in the presence (black) and absence (red) of 1mM DTT. Bottom: The same comparison for 155 nM EcUvrD(V425C). **(C)** The fraction DNA unwound by EcUvrD(R421C) is shown as a function of time for four conditions: (1) In excess DNA +DTT (green), in excess DNA –DTT (cyan), in excess protein +DTT (black), and in excess protein –DTT (red). **(D)** The fraction DNA unwound by EcUvrD(V425C) is shown in the same conditions.

Single-round fluorescence stopped-flow kinetics experiments were performed with a 3’-(dT)_20_-18 bp DNA substrate as described(*32, 39*) to assess the activity of the mutated *E. coli* construct (Supplemental Methods). No helicase activity is observed under conditions that favor monomers bound to DNA (*i*.*e*., excess DNA (25 nM) over protein (12.5. nM) under reducing conditions (+DTT, Fig. 5C, green). However, robust helicase activity is observed at the same concentrations under oxidizing conditions (–DTT), indicating that helicase activity correlates with the formation of the disulfide linked *Ec*UvrD(R421C) dimers (Fig. 5C, cyan). When *Ec*UvrD(R421C) (25 nM) is in excess over DNA (12.5 nM), helicase activity was observed even in the presence of DTT indicating that *Ec*UvrD(R421C) can still form non-crosslinked dimers (Fig. 5C, black). This activity was also stimulated substantially in the absence of DTT due to the additional formation of crosslinked dimers (Fig. 5C, red).

In contrast, a different cysteine mutation (V425C) that was made based only on a sequence alignment with UvrD1 does not display redox-dependent dimerization (Fig. 5B, bottom panel) nor oxidative-dependent helicase activation (Fig. 5D). These data indicate that the same 2B subdomain dimerization interface observed in the structures of *Mt*UvrD1 is conserved in *Ec*UvrD, suggesting that the 2B dimerization mechanism is a general feature of UvrD-family enzymes.

## Discussion

We report the first structures of a dimeric UvrD-family SF1A helicase obtained in solution conditions that support helicase activity (*32*). The *Mt*UvrD1 dimer structure reveals two novel aspects of these enzymes. Firstly, in contrast to the *Ec*UvrD and *Bs*PcrA monomeric DNA-bound structures (*19, 20*), the 2B domain does not contact duplex DNA but forms the dimer interface. Secondly, 2B subdomain flexibility allows the two subunits of the dimer to arrange their motor domains so that they are oriented in the same direction on the ssDNA, consistent with its 3’ to 5’ translocation directionality. This flexibility also allows for a path to be opened in the leading subunit to allow ssDNA to pass to the trailing subunit.

These structures highlight the critical regulatory role of the 2B subdomain in activation and regulation of helicase activity by dimerization. Residues that contact the DNA in monomeric complexes (*19, 20*), including the GIG motif, are instead found far from DNA and close to the dimerization interface of UvrD1 (Fig. 4C). Hence 2B dimerization and duplex DNA binding by the 2B domain are competitive leading to a straightforward model for helicase activation: Dimerization results in a major reorientation of the 2B subdomain so that it is unable to interact with the duplex DNA, thus relieving its auto-inhibition as suggested previously (movie S9) (*3, 34*). This also means that the monomer-DNA structures need to be reinterpreted as inhibited complexes rather than the active helicase conformations. Our ability to create a disulfide bonded *Ec*UvrD dimer based on the *Mt*UvrD1 structures further suggests that dimerization via 2B subdomains is likely a general mechanism for UvrD-family helicase activation. In fact, the 2B subdomains of RecB and RecC form a dimerization interface in RecBCD and do not interact with DNA, although the 2B-2B interface differs from that observed in the UvrD1 dimer (*40*).

Despite robust biochemical evidence to the contrary, there has been debate concerning whether UvrD-family enzyme monomers possess processive helicase activity. The claim that Rep, PcrA and UvrD monomers are active helicases is based primarily on the existence of structures of monomers of *Bs*PcrA (*19*) and *Ec*UvrD (*20*) bound to DNA junctions coupled with the fact that monomers are able to translocate directionally along ssDNA (*22-24, 34*). The presence of monomeric crystal structures that appear poised to unwind DNA coupled with the absence of dimeric structures has had an overwhelming influence in the field.

In particular, the observation that the 2B subdomain in the monomeric structures interacts with duplex DNA led to the hypothesis that this interaction is essential for DNA unwinding (*19, 20*). However, UvrD, PcrA and Rep monomers are not active helicases on their own. Monomers can processively translocate directionally along ssDNA coupled to ATP binding and hydrolysis, but ssDNA translocation is not sufficient for helicase activity (*3, 23-25, 27, 41*). In the absence of a destabilizing force on the DNA, UvrD-family monomers need to be activated. Activation can occur by dimerization (*26, 27, 29, 31-33, 42*), or by interactions with accessory proteins: PriC for Rep (*43*), MutL for UvrD (*39, 44*), RepD for PcrA (*45*). In the case of Rep, deletion of the 2B subdomain actually activates the helicase activity of the monomer indicating that the 2B subdomain of Rep is auto-inhibitory and not needed for DNA unwinding (*28, 34*). In addition, activation via UvrD-MutL interactions is accompanied by rotation of the 2B subdomain to an intermediate conformation (*44*). Previous mutations made in the 2B subdomain to “test” whether disruption of the 2B-DNA contacts in the monomer-DNA structures are catalytically important did result in reduced helicase activity (*20*). However, based on the UvrD1 dimer structures presented here, those same mutations would lead to reduced helicase activity due to dimer destabilization. It is worth noting that R421, which we mutated to Cys in *Ec*UvrD to form active disulfide linked *Ec*UvrD dimers, directly interacts with duplex DNA in the *Ec*UvrD monomer-DNA structure (*20*). This is consistent with the fact that the monomer structure does not represent the active helicase since the R421-DNA interaction cannot occur in the UvrD dimer-DNA complex. The claim that a UvrD monomer-DNA crystal structure captures the melting of one base pair by the helicase is not justified as the complex was formed using a DNA substrate with a “pre-melted”, unpaired T–T mismatch at the ssDNA–dsDNA junction (*20*).

We note that monomers of Rep can display helicase activity when the 2B subdomain is crosslinked intramolecularly in a relatively closed conformation (Rep-X) (*46*). UvrD monomers can also display limited helicase activity when a destabilizing force is applied to the DNA duplex (*47, 48*). The mechanism by which intramolecular crosslinking of the 2B subdomain activates the monomer helicase is not understood but it may constrain it to a position where it cannot rotate to interact with duplex DNA, consistent with the model we propose for activation. In fact, intramolecular crosslinking of the 2B subdomain of Rep in a relatively open conformation (Rep-Y) that should prevent it from interacting with duplex DNA does not eliminate helicase activity (*46*). The activation via force is more straightforward as it tilts the free energy landscape towards unwound DNA and lowers the energy barrier for DNA unwinding.

Although the monomeric UvrD and PcrA structures do not represent conformations of active helicases, we suggest that those structures may explain a mechanism to prevent unwanted helicase activity. In addition to its role as a helicase, UvrD also acts as an anti-recombinase by using its monomeric ssDNA translocase activity to displace RecA protein filaments from ssDNA (*49*). Once UvrD removes a RecA filament from a ssDNA gap, it will encounter a 3’-ss/ds DNA junction. In order to prevent unwanted helicase activity, the 2B subdomain of the monomer binds to the duplex DNA to inhibit DNA unwinding.

What might be the biological roles for a UvrD dimer? We note that *Ec*UvrD is also involved in transcription-coupled DNA repair (TCR). A UvrD monomer is proposed to be continuously bound to RNA polymerase (RNAP) during normal growth conditions (*15*). However, upon UV irradiation, a second UvrD monomer binds RNAP to form a UvrD dimer that facilitates backtracking of RNAP during TCR. Crosslinking mass spec studies suggest that the UvrD dimer interface involves the 2B subdomains of each subunit (*15*), consistent with the *Mt*UvrD1 dimer structure presented here.

The structures reported here provide insight into how SF1A helicases are activated and regulated by the 2B subdomain. It is known that both subunits of the functional dimer need to hydrolyze ATP (*27, 33*) and that helicase activity involves allosteric communication among the ATP sites, the DNA binding sites and the dimer interface (*27, 33, 50-52*). As previously suggested (*18*) and shown here, the inherent flexibility allowed by rotation about the hinge connecting each 2B subdomain to the rest of the subunit enables the dimer to undergo large conformational changes that are likely involved in DNA translocation and unwinding mechanisms (*27, 31, 33, 50, 53*). How each subunit moves relative to one another during translocation and DNA unwinding remains to be determined. Further biochemical and biophysical studies as well as additional structures of intermediates along the pathway will be required to understand the mechanistic details of DNA unwinding by these ubiquitous enzymes. However, our identification of the 2B subdomain as a general dimerization motif for SF1A helicases, coupled with the dependence of helicase activity on dimerization, identifies the subdomain as a crucial regulatory module for the activation or repression of DNA unwinding by these enzymes.

## Acknowledgements

We thank the staff of the Washington University Center for Cellular Imaging including Drs. Katherine Basore, Brock Summers, and Brad Readnour for their indispensable support in the acquisition and processing of the cryoEM data. We also acknowledge Drs. Eric Tomko and Kacey Mersch for fruitful discussions and Drs. Weikai Li, Rui Zhang, and Alireza Ghanbarpour for their assistance during structural modeling.

## Author contributions

Conceptualization: AC, TML, EAG

Methodology: AC, BN, TML, EAG

Investigation: AC, BN

Visualization: AC, BN, TML, EAG

Funding Acquisition: TML, EAG

Project administration: TML, EAG

Writing: AC, TML, EAG

## Data availability

The EM maps are available on the Electron Microscopy Database as follows: apo compact (EMD-46752); apo extended (EMD-46753); DNA-bound dimer (EMD-46797 and EMD-46799); DNA-bound monomer (EMD-46850). The molecular models are available in the PDB as follows: apo compact (9DCI); DNA-bound dimer (9DES); DNA-bound monomer (9DGY).

## Supplementary Materials

Materials and Methods

Figures S1 to S14

Table S1

References (1-16)

Movies S1 to S9

