## Supplemental Information for "Structural Basis for Dimerization and Activation of UvrD-family Helicases"

Ankita Chadda[1], Binh Nguyen[1], Timothy M. Lohman[1] and Eric A. Galburt[1]
[1]Department of Biochemistry and Molecular Biophysics, Washington University in Saint Louis School of Medicine,
Saint Louis, MO 63110

#### Materials and Methods

**Buffers and Reagents: Buffers and Reagents:** Buffer G is 20 mM Tris pH 8.3, 20% (v/v) glycerol, 1.0 mM EDTA, 0.5mM EGTA, 5 mM 2-mercapto-ethanol, varying NaCl. Buffer T is 10 mM Tris pH 8.3, 20 mM NaCl, 20% (v/v) glycerol. Stocks of disodium ATP solution (Sigma-Aldrich) in pH 7.5 as described (*1*). Stock solution of $MgCl_2$ were prepared in Milli-Q water, and its concentration was determined using refractive index measurements as described (*1*).

**Protein purification:** *Mt*UvrD1 both wild type and the 2B cysteine to alanine point mutant at position 451 were expressed and purified as described (*2*) except that Triton X100 was removed from the buffers after Ni-NTA elution. Ion exchange chromatography, size exclusion, and protein storage was performed in buffers lacking detergent. The protein was stable at -80 °C for up to 6 months.

The *E. coli* UvrD(R421C) and UvrD(V425C) mutations were introduced using a QuikChange kit (Stratagene, Cedar Creek, TX) into a plasmid (pGG209ΔCys) encoding a Cys-less UvrD gene in which all six naturally occurring Cys had been mutated to Ser (*3*). *E. coli* BL21(DE3)ΔUvrD cells were transformed with the plasmid and grown in Terrific Broth with 50ug/ml Kanamycin at 37 °C and 220 rpm. At $OD_{600}$=0.8, the cultures were chilled on ice for 30 minutes, IPTG was added to 0.3 mM final and grown overnight with shaking at 16 °C. Mutations were confirmed by DNA sequencing. UvrD(R421C) and UvrD(V425C) proteins were purified as described (*4*) with the following modifications. In brief, the clear lysate containing the 6xHis-tagged UvrD mutant was loaded onto HisPur™ Ni-NTA (Thermo Scientific) and washed, and the targeted protein was eluted with buffer G containing an imidazole gradient (10-250 mM). The pooled fractions were diluted with buffer G to 0.2 M NaCl then loaded onto HiTrap heparin HP (Cytiva), and the protein was eluted with buffer G containing 0.2 to1.0 M NaCl gradient over 20 column volumes). The pooled fractions were diluted with buffer G to 0.15 M NaCl and loaded onto a tandem dsDNA -ssDNA column connected in series at 75 mM NaCl. The UvrD mutants passed through the dsDNA and bound to the ssDNA column. After disconnecting the columns, the ssDNA column was washed with buffer G + 0.2 M NaCl, and the UvrD mutant was eluted with buffer G+ 2.0 M NaCl. The 6xHis-tag was removed by thrombin digestion (Sigma-Aldrich) (Buffer G + 0.2M NaCl, 2 NIH units per mg of tagged-protein for 24 hours at 4 °C). The un-cleaved and cleaved proteins were separated by passing through the Ni-NTA column. The thrombin was removed with the use of the ssDNA column or the HiTrap Benzamidine FF column (Cytiva). UvrD mutant concentrations were determined using an extinction coefficient $\varepsilon_{280}=1.06\times10^5$ $M^{-1}$ $cm^{-1}$.

**Crosslinked *Ec*UvrD(R421C) dimers:** Disulfide crosslinked *Ec*UvrD(421C) dimers were prepared as follows. *Ec*UvrD(R421C) protein at 9.5 µM monomer (in 20 mM Tris-HCl pH 8.3, 0.2 M NaCl, 1 mM EDTA, 50% (v/v) glycerol, 5 mM 2-mercaptoethanol) was dialyzed vs. buffer at lower [NaCl] and without reducing agent (10 mM Tris-HCl pH 8.3, 20 mM NaCl, 50 % (v/v) glycerol) to remove the 2-mercaptoethanol. The final protein concentration after dialysis was 6.90 µM. The DNA unwinding and AUC experiments were performed in buffer T with this stock concentration.

**DNA substrates:** DNA for the Cryo-EM experiments was formed by annealing two oligomers purchased from IDT purified by PAGE. Specifically, a 38 nt single-stranded DNA (5'-GTT GGT CGG CAG CAG GGC $(T)_{18}$-3') was mixed with an equimolar concentration of a complementary 18 nt DNA strand (5'-GCC CTG CTG CCG ACC AAC-3') in 10 mM Tris, pH 8.0, and 50 mM NaCl, followed by heating to 95 °C for 5 min and slow cooling to room temperature resulting in a partial duplex of 18 base-pair with a 3'-$dT_{20}$ ssDNA tail. The DNA substrate used for the stopped-flow helicase experiments were purchased from IDT DNA with HPLC purification (Coralville, IA, USA). The extinction coefficients (per strand) are: (for Cy5-strand: 5'-Cy5-GTT GGT CGG CAG CAG GGC $(T)_{20}$-3' ($\varepsilon_{260,dna}$=3.32×10$^5$ M$^{-1}$ cm$^{-1}$) and for BHQ2-strand: 5'-GCC CTG CTG CCG ACC AAC-BHQ2-3' ($\varepsilon_{260,dna}$=1.57×10$^5$ M$^{-1}$ cm$^{-1}$)). Other extinction coefficients are: Cy5 ($\varepsilon_{260}$=1.0×10$^4$ M$^{-1}$ cm$^{-1}$, $\varepsilon_{650}$=2.5×10$^5$ M$^{-1}$ cm$^{-1}$) and BHQ2 ($\varepsilon_{260}$=8.0×10$^3$ M$^{-1}$ cm$^{-1}$, $\varepsilon_{579}$=3.8×10$^4$ M$^{-1}$ cm$^{-1}$). The trap DNA is 5'-GCC TCG CTG C-$(T)_5$-G CAG CGA GGC-$(T)_{40}$-3' ($\varepsilon_{260}$=5.41×10$^5$ M$^{-1}$ cm$^{-1}$).

**Stopped-flow DNA helicase assays:** DNA helicase activity of *Ec*UvrD(R421C) and *Ec*UvrD(V425C) was examined at 25 °C buffer T (± 1 mM dithiothreitol (DTT)) under single round conditions using an Applied Photophysics SX.18MV stopped-flow instrument (Applied Photophysics Ltd., Leatherhead, UK) as described (*5*). The DNA substrate was an 18 bp duplex DNA containing a 3'-$(dT)_{20}$ tail with a Cy5 fluorophore attached to the long strand and Black Hole Quencher 2 (BHQ2) attached to the short strand. When in close proximity, BHQ2 quenches Cy5 fluorescence, hence, DNA strand separation is accompanied by an increase in Cy5 fluorescence. DNA unwinding was initiated by mixing pre-formed UvrD-DNA complexes with buffer T containing 1 mM ATP, 2 mM $MgCl_2$ and 2 µM protein trap. The protein trap, a 10 bp DNA hairpin (DNA X) possessing a 3'-$(dT)_{40}$ ssDNA tail, binds to any free UvrD or UvrD that dissociates during unwinding thus preventing any re-initiation of DNA unwinding and ensuring that any unwinding is due to UvrD that is pre-bound to the DNA (*i.e.*, single-round conditions). The concentrations after mixing were 25 nM protein + 12.5 nM DNA (protein > DNA; dimer conditions) or 12.5 nM protein + 25 nM DNA (protein < DNA; monomer conditions). All reported traces are the average of 15-20 replicate stopped flow shots.

**Analytical Sedimentation:** Sedimentation velocity experiments were performed at 25 ℃ in Buffer T (±1 mM DTT) at 42,000 rpm using a Proteome Lab XL-A analytical ultracentrifuge equipped with an An50Ti rotor as described (*2*). Crosslinked EcUvrD(R421C) protein samples (155 nM and 175 nM), prepared as described above, were scanned at 230 nm and data analyzed using SEDFIT to obtain sedimentation coefficient distributions (*6*). Sedimentation coefficients were corrected to $s_{20,w}$ as described (*2*).

**CryoEM sample preparation:** For apo UvrD1 complexes, purified *Mt*UvrD1 was concentrated to 20 µM and 400 µl was dialyzed in 1 L TRIS at pH 8.0 in addition to 75 mM NaCl and 5% glycerol and 6 µM UvrD1 (monomer units) was crosslinked in the presence of 0.5 mM BS3 crosslinker (Thermo Fisher Scientific, #P121580) at room temperature for 45 minutes. Subsequently, 30 mM TRIS at pH 7.5 was added for 15 minutes to quench unreacted crosslinker. Crosslinked protein was dialyzed in 1 L phosphate buffer at pH 8.0, 5% glycerol and 75 mM NaCl for 2-3 hours at room temperature to remove residual crosslinker and then immediately used to prepare cryoEM grids.

For UvrD1-DNA complexes, purified *Mt*UvrD1 was concentrated to 20 μM and 400 μl was dialyzed in 1 L phosphate buffer a 1:1 molar mixture of UvrD1 and 3'-d$T_{20}$-18 bp partial duplex was used in the presence of AMPPNP and $MgCl_2$ (6 μM UvrD1 (monomer units), 6 μM 3'-d$T_{20}$-18bp partial duplex, 1 mM AMPPNP, and 5 mM $MgCl_2$). Excess DNA was used to saturate the protein and facilitate the capture of protein-DNA particles due to the superior contrast of the protein. BS3 crosslinker was added to the above reaction mixture at room temperature to about 45 minutes. Next 30 mM TRIS, pH 7.5, was added to the reaction mixture for an additional 15 minutes to quench the unreacted crosslinker. The sample was dialyzed for 2-3 hours at room temperature with phosphate buffer at 5% glycerol and 75 mM NaCl to remove residual crosslinker and immediately used to prepare cryoEM grids.

Grid Freezing and Imaging: For each sample, the Vitrobot Mark IV (ThermoFisher) was used to freeze grids with 6 μM UvrD1 (monomer units). Samples of UvrD1 dimer alone and in complex with DNA were imaged on a ThermoFisher Glacios microscope and the UvrD1 monomer-DNA complex was imaged on a ThermoFisher Krios microscope. Imaging parameters are provided in Supplementary Table 1.

**CryoEM data processing:**

Apo *Mt*UvrD1 (Fig. S1-3): For data processing of apo UvrD1, 4562 images were acquired in which 2680 movies were at normal acquisition and 1882 were acquired at tilts of 10- and 20-degrees. Image analysis was performed using cryoSPARC software (*7*). Images were subjected to patch motion correction for correcting global and local motion and patch-based CTF to automatically detect defocus variation for tilted and bent samples. Low-quality integrated movies with a CTF lower than 5 Å were excluded using manual exposure tools. Particles were picked using blob picker with particle dimensions of 70 Å or 200 Å for both monomer and dimer respectively resulting in 2.3 million particles. Particles were then extracted using box size of 350 Å and Fourier cropping at 200 Å (increasing the pixel size from 0.9 to 1.6). After extraction, 1.5 million particles were used for first round of 2D classification into 200 classes that gave 1-2 good classes with defined features that were used as templates to pick particles and perform another 3 rounds of template picking, extraction, and 2D classification using the above parameters. After the 3$^{rd}$ round of 2D classification, only the best classes with well-defined features (175,329 particles) were selected to generate initial reference-free ab-initio models. This resulted in four classes: A junk class, one low-resolution monomer class, and two dimer classes with almost equal number of particles (50,845 and 46,812) representing the compact and extended dimer forms respectively. The dimer classes were further refined using heterogenous, non-uniform, and local refinement resulting in 3D volumes of UvrD1 compact and extended conformations at 5.5 Å and 6.6 Å resolution.

For further analysis of the compact apo conformation, an additional 1653 movies were collected bringing the total 6213. Templates were created from the 5.5 Å volume obtained above and 2D classification increased the number of good particles to 697,920 that were used for ab initio modeling. From reference free modeling into 4 models, one of them with 189,910 particles looked like the apo-compact dimer that was further subjected to heterogenous and non-uniform 3D refinements to get a 3D model at 4.63 Å resolution. To resolve flexibility and improve resolution further, 189,910 particles were subjected to multiple rounds of 2D classification and 3D non-uniform refinement using C2 symmetry to get a final model of compact form of apo UvrD1 at a 4.07 Å global resolution with local resolution range from 3.6 to 6.7 Å where exposed helices are at a lower resolution than the protein interior. For processing of extended conformation of apo UvrD1, 3D model obtained using 50,845 particles above were used to create templates and perform another round of particle picking and 2D classification from which 369,486 particles were selected for reference free ab-initio modeling into 4 models where 109,470 particles belonged to the extended confirmation. After multiple rounds of 2D classification of these 109,470 particles and ab-initio modeling with 3D refinement gave a model of the extended conformation of apo

UvrD1 at 4.84 Å global resolution with local resolution range being 4.6 to 8 Å (EMD entry ID: EMD-46752). The extended conformation was refined using C1 symmetry (EMD entry ID: EMD-46753)

<u>*Mt*UvrD1 dimer bound to DNA (Fig. S6-10):</u> UvrD1 (6 μM monomer units) was incubated with 3'-dT$_{20}$-18bp DNA (6 μM), 1 mM AMPPNP and 5 mM $MgCl_2$ prior to freezing and image collection. 2211 movies were acquired using Glacios 200 kV. Images were subjected to patch motion correction for correcting global and local motion and patch-based CTF to automatically detect defocus variation. Low-quality integrated movies with a CTF lower than 5 Å were excluded using manual exposure tools. After performing blob picking and initial 2D classification good looking classes were used as templates for template picking and particles were extracted using 350 Å box size with Fourier cropping at 200 Å (increasing the pixel size from 0.9 to 1.6). Multiple rounds of 2D classifications gave good 2D classes with 493,445 particles that were used for a reference free ab initio and heterogenous refinement that gave five classes with one class of 34% DNA bound to UvrD1 dimer and one class 23% apo dimer conformation. The other three classes did not give a define 3D volume and hence were classified as junk. The DNA bound UvrD1 density was at low resolution and had considerable heterogeneity because of the flexibility of the 18 base-pair DNA. The DNA bound conformation was next subjected to homo refinement and to resolve the issue of heterogeneity and to visualize double stranded DNA, two input volume were provided, one volume including two subunits and one including subunits and the DNA that improved the resolution of double stranded DNA bound structure at an overall global resolution of 5.9 Å (EMD entry ID: EMD-46797). For visualizing single stranded DNA threading through both subunits additional 2336 images were acquired with total being 4547 that were again subjected to Patch motion correction and patch CTF and particle picking using blob picker with 70-200 Å blob dimension settings which picked around 4.3 million particles that were subject to extraction and 2D classification to give good 2D classes that were used as templates for template picking. Performing multiple rounds of 2D classification gave good classes with 634,557 particles that were subjected to ab initio and heterogenous refinement into five classes. In this refinement 34% were DNA bound UvrD1 dimer and 23% particles in apo dimer conformations. For visualizing single stranded DNA within the two subunits, the DNA bound structure (34%) with 209,170 particles was first subjected to non-uniform refinement and then 3D classification with four classes providing a focused mask around both the subunits of a dimer that improved the resolution of one class with 51,008 particles to a resolution of 4.9 Å with local resolution range between 3.5-6.5 Å (EMD entry ID: EMD-46799). All 3D refinement for DNA-bound structures were performed using C1 symmetry.

<u>UvrD1 monomer bound to DNA (Fig. S15-16)</u>: For analyzing UvrD1 monomer bound to DNA, 6 μM of the 2B cysteine mutant (C451A) was incubated with 6 μM of 3'-dT10-18bp DNA in the presence of 1 mM AMPPNP, 5 mM $MgCl_2$ prior to freezing. 4999 movies were acquired on Krios microscope. The images were subjected to patch motion correction and CTF. Particle picking using blob picker with 70-120 Å dimension settings picked around 5.1 million particles that were subject to extraction using box size 250 Å and 2D classification to give few 2D classes that were used as templates for template picking. Performing 3 rounds of particle picking using templates got 713,562 particles that were subjected to reference free ab initial modeling and heterogenous refinement with 4 models in which two classes (total particle number 407,020) represented the DNA bound UvrD1 conformation. These two classes when refined separately via non-uniform refinement but did not provide a complete 3D model. The particles belonging to the above two 3D refinement jobs were thus re-extracted with a box size of 280 Å and went another round of 2D classification out of which 282,380 were selected and went into 3 class ab-initio and then heterogenous refinement. One of the classes was further refined using non-uniform refinement that gave the low-resolution density map of monomer bound to DNA at 5.61 Å (EMD entry ID: EMD-46850). No symmetry was applied during the refinement procedures.

Modeling and Refinement: The predicted structure of monomer of *Mt*UvrD1 was obtained using PHYRE2 software (*8*) by submitting its amino acid sequence. The PDB reference used for modeling previously observed open and closed structures of *Mt*UvrD1 was *Ec*UvrD 3LFU (*3*), *B. stearothermophilus* PcrA helicase complex 3PJR (*9*) and *Ec*UvrD 2IS4 (*10*). The 1A, 1B, 2A and 2B subdomains for the UvrD1 model were assigned using the structural alignment of the *Ec*UvrD monomer with the homology model of *Mt*UvrD1. The model for the apo UvrD1 dimer in the compact conformation was built in ChimeraX (*11*) by manually docking the two open conformation homology models into the density and then using the Volume Fit tool with a resulting map-to-model correlation coefficient of 0.85. This was further subjected to rounds of modeling in COOT (*12*) and refinement in Phenix (*13*). Since the homology model was built in the absence of the C-terminal Tudor domains, these were docked manually in Chimera using models derived from AlphaFold . This model of the compact apo structure has PDB ID code 9DCI.

For the DNA bound dimer structure, 18 base-pairs of DNA duplex with 3'-$dT_4$ single stranded from *D. radiodurans* UvrD 4C2T (*14*) was modeled into the leading subunit density and 3'-$dT_{16}$ ssDNA and *E. coli* Rep helicase DNA complex 1UAA (*15*) was docked into density spanning the motor domains of the two subunits. Modeling and refinement resulted in 10 single-stranded nucleotides fit in density. Multiple rounds of building in COOT and refinement in Phenix using both the masked and unmasked maps led to the final model (PDB ID: 9DES).

For the DNA bound monomer structure, the model of a monomer of *Ec*UvrD bound to DNA (*10*) was first fit into the density using ChimeraX Volume Fitting. A model of *Mt*UvrD1 was aligned with the *E. coli* protein. The sequence of the DNA model was adjusted to match the DNA substrate used in our experiment. The resulting *Mt*UvrD1-DNA complex went through multiple cycles of modeling in COOT and refinement with Phenix leading to the final model (PDB ID: 9DGY).

### Figures S1 to S12:

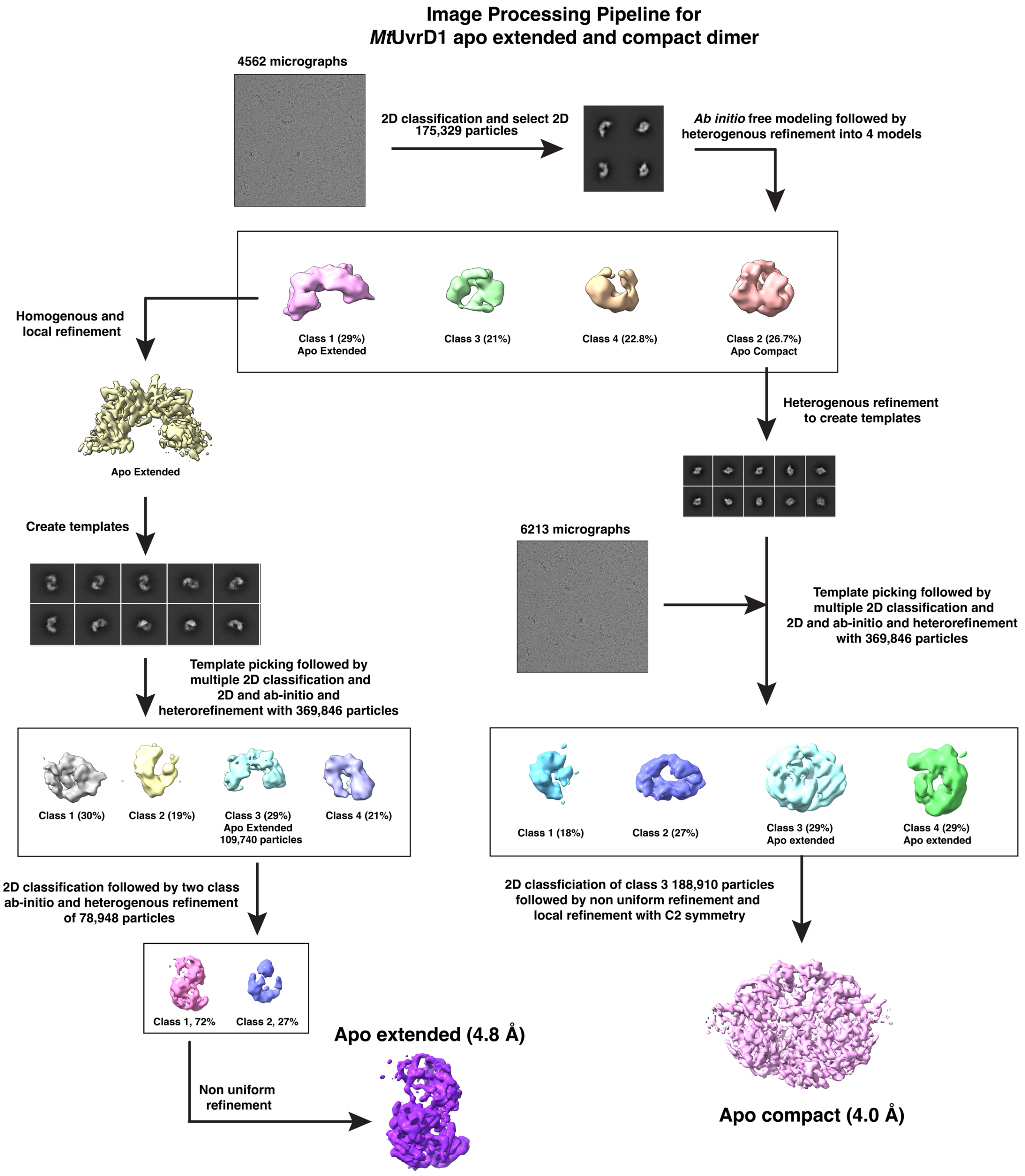


**Fig. S1:** Image processing pipeline for *Mt*UvrD1 apo compact and extended maps.

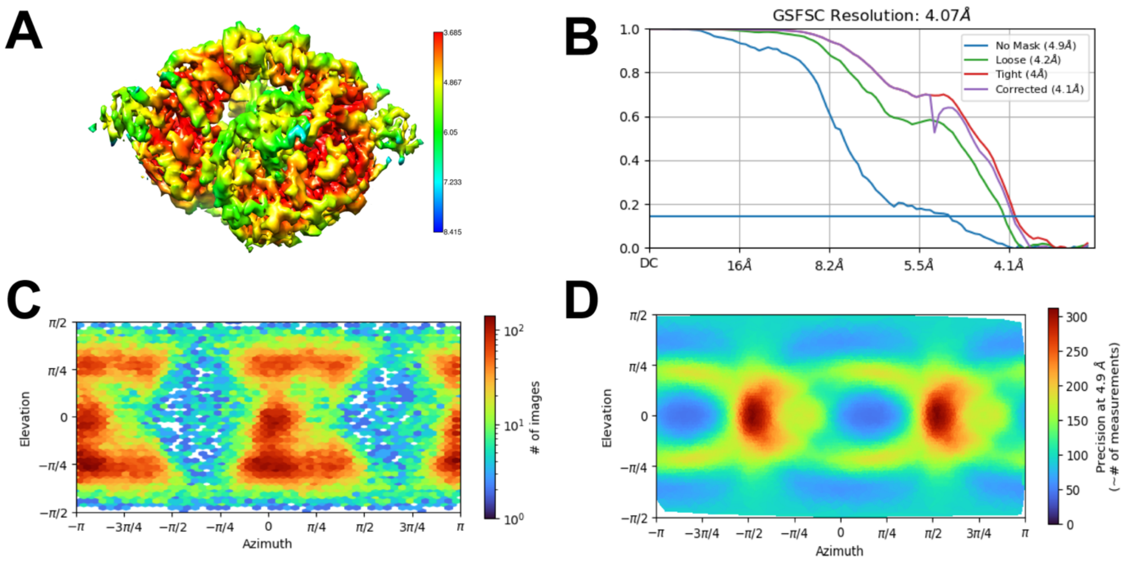


**Fig. S2:** Statistics for apo UvrD1 compact dimer. **(A)** Local resolution. **(B)** FSC curves. **(C)** Particle orientation distribution, and **(D)** Posterior directional distribution of particles.

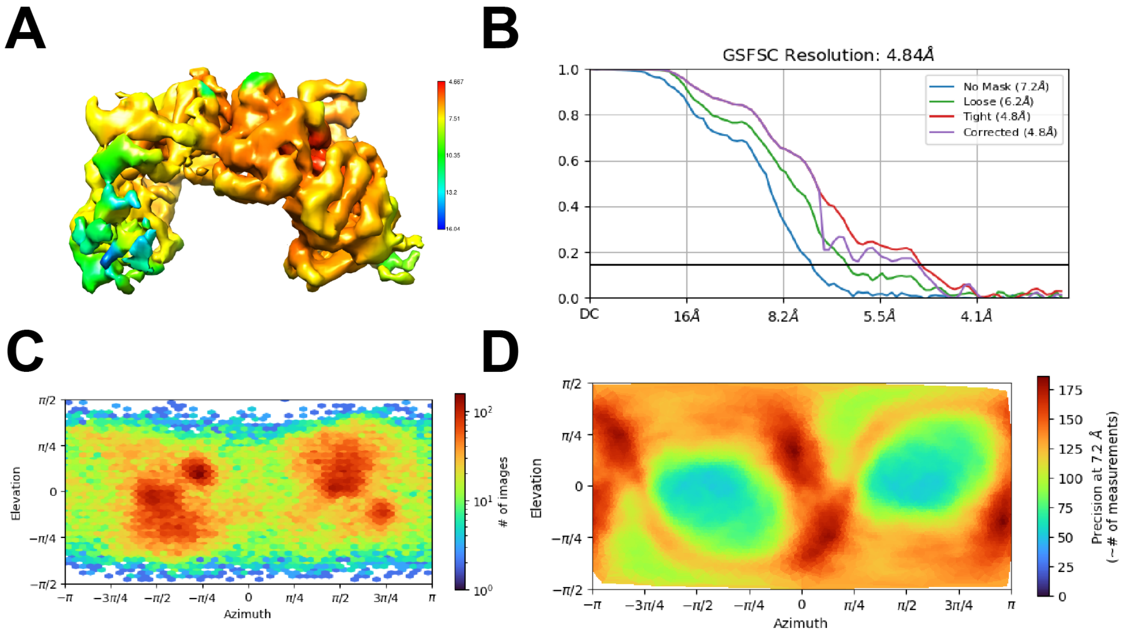


**Fig. S3:** Statistics for apoUvrD1 extended dimer. **(A)** Local resolution. **(B)** FSC curves. **(C)** Particle orientation distribution, and **(D)** Posterior directional distribution of particles.

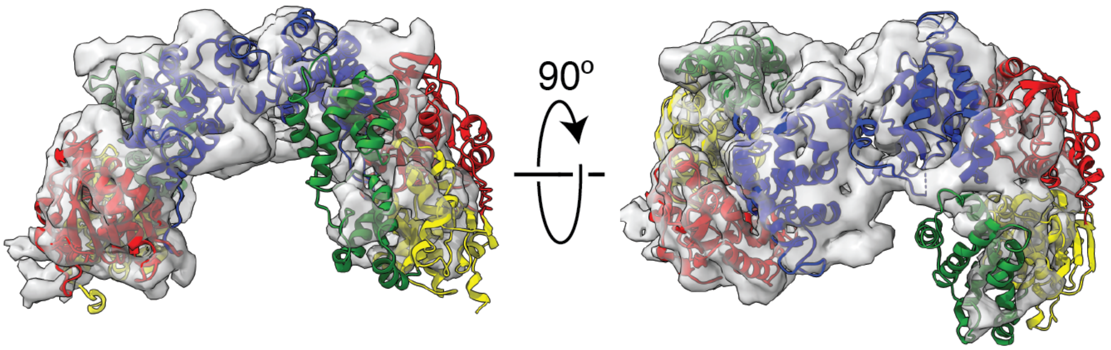


**Fig. S4:** Best fit of *Mt*UvrD1 dimer into extended conformation density. The left subunit density is better resolved than the right subunit suggesting a fair amount of conformational heterogeneity around the dimer interface. The domains are colored as in the text: 1A (yellow), 1B (green), 2A (red), 2B (blue).

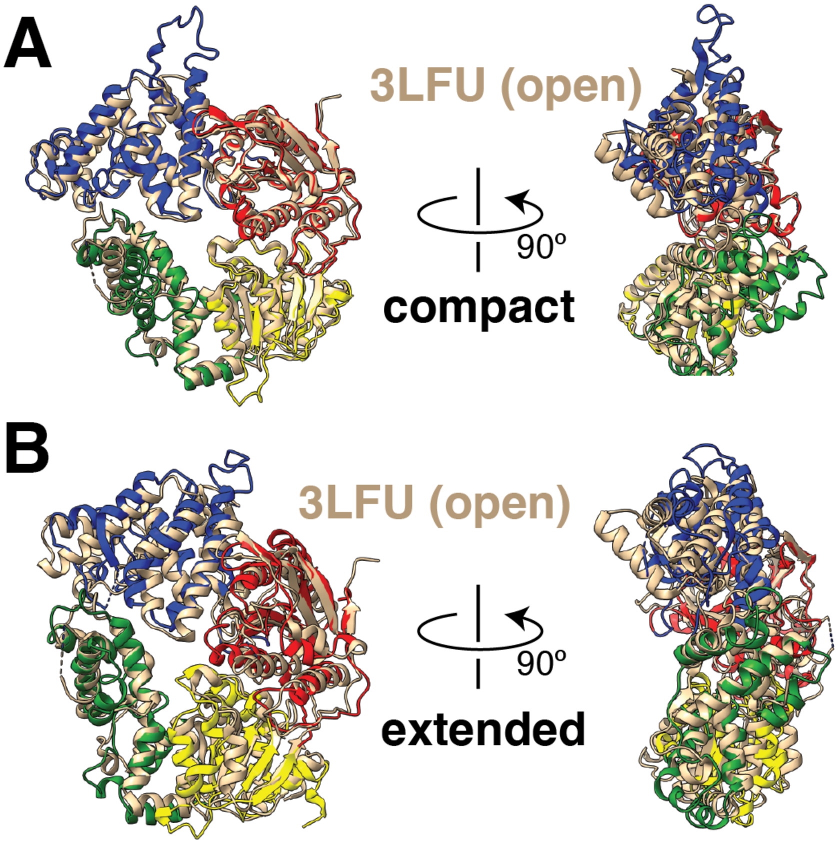


**Fig. S5:** Overlay of EcUvrD open structure (3LFU, tan) and compact apo *Mt*UvrD1 color coded by domain.

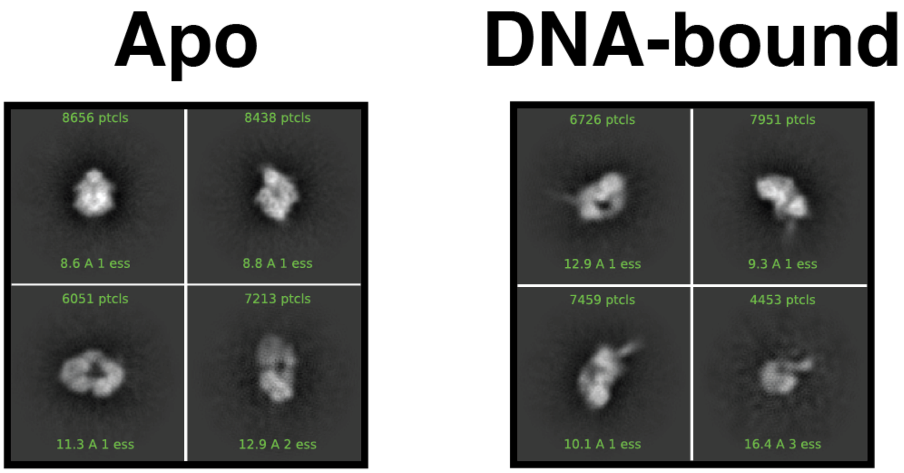


**Fig. S6:** 2D classes of dimer-DNA bound sample showing the existence of apo-compact and DNA-bound particles.

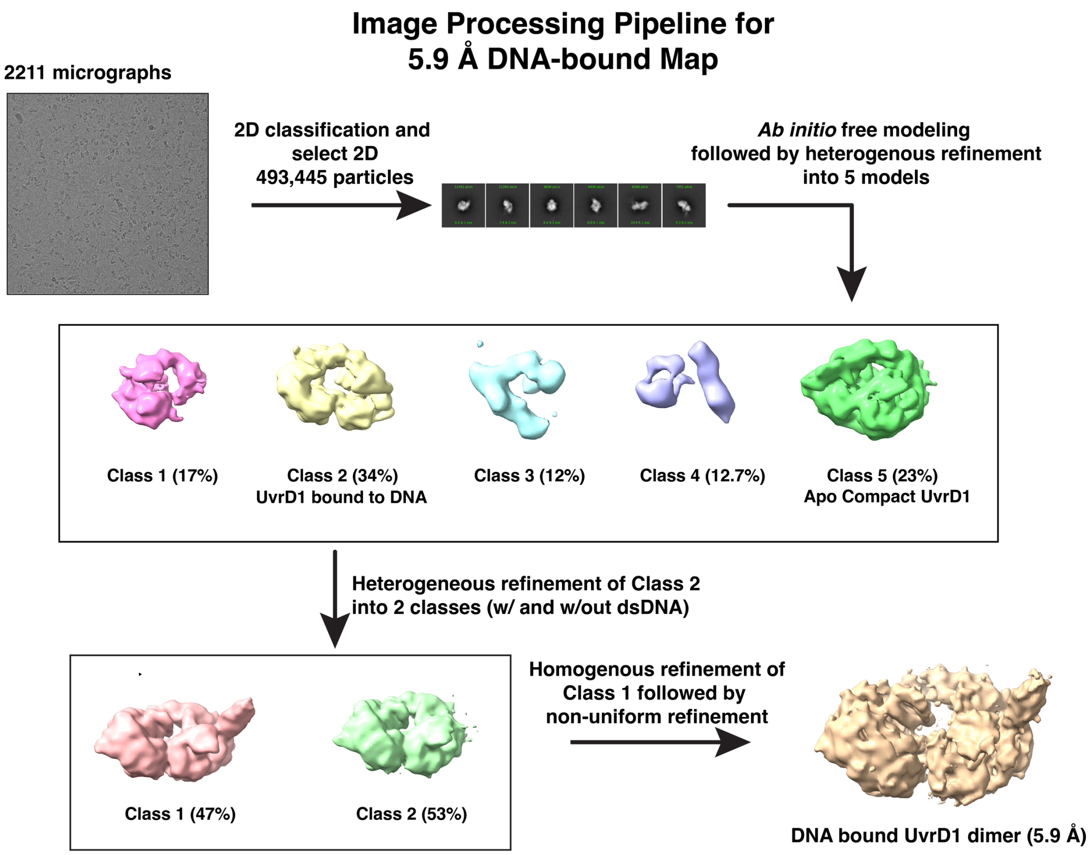


**Fig. S7:** Image processing pipeline for 5.9 Å DNA-bound *Mt*UvrD1 map.

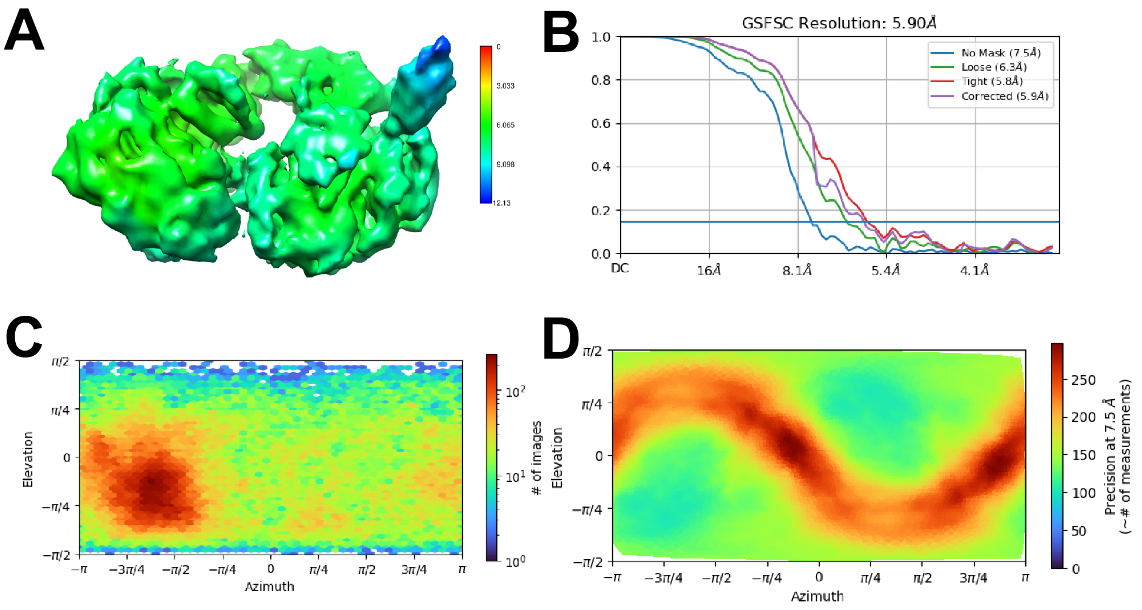


**Fig. S8:** Statistics for initial dimer complex. **(A)** Local resolution. **(B)** FSC curves. **(C)** Particle orientation distribution, and **(D)** Posterior directional distribution of particles.

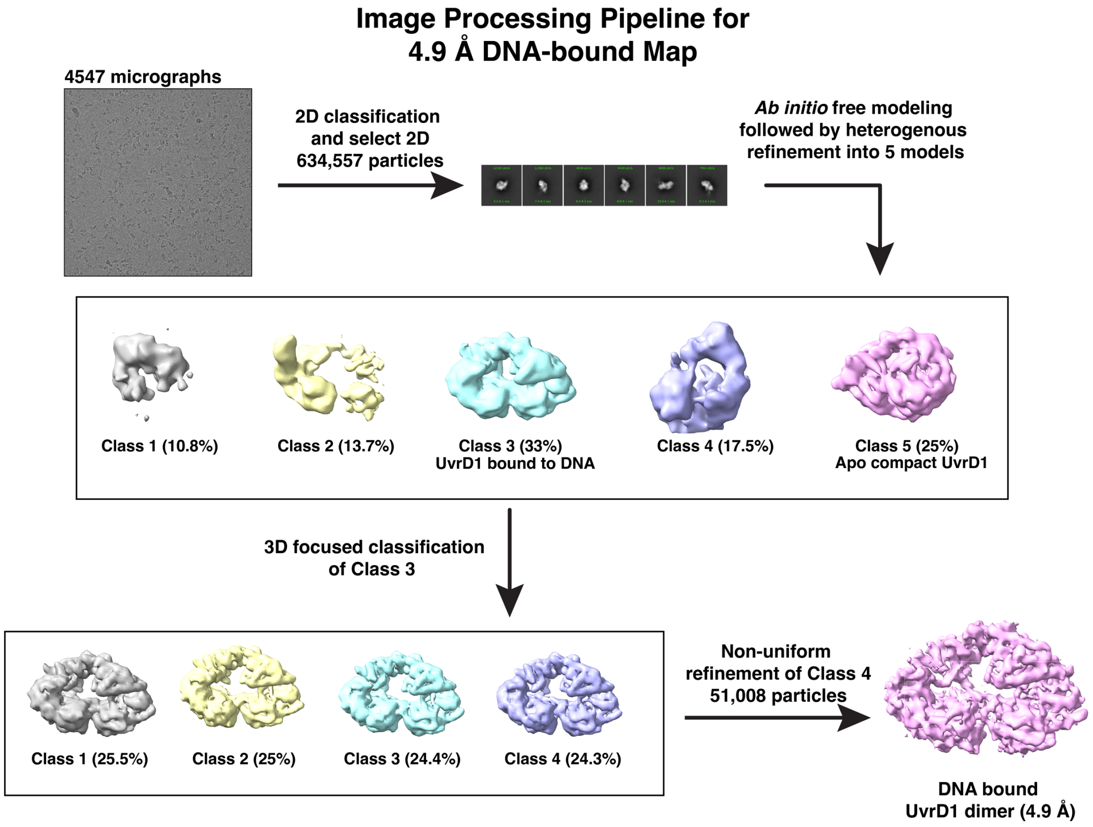


**Fig. S9:** Image processing pipeline for 5.9 Å DNA-bound *Mt*UvrD1 map.

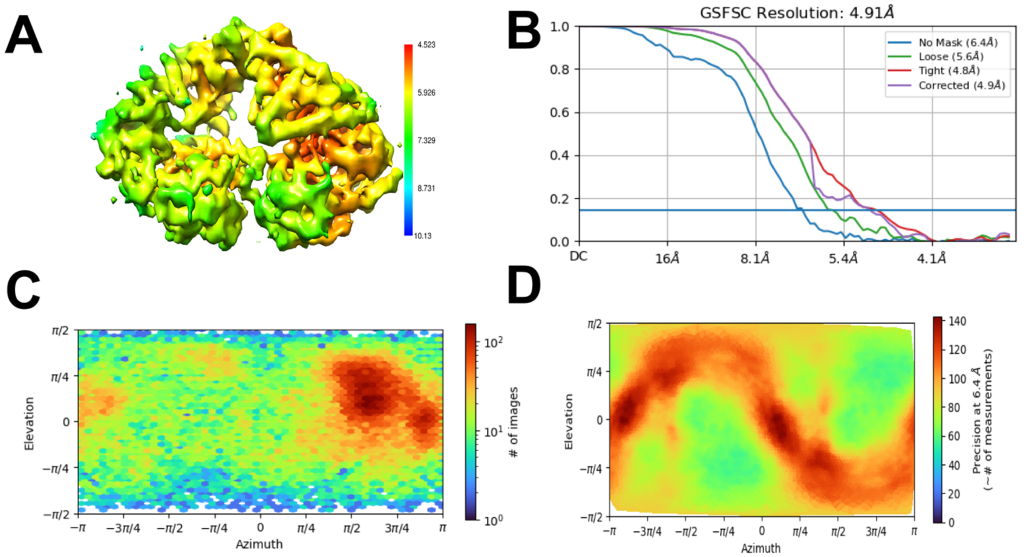


**Fig. S10:** Statistics for masked DNA-bound dimer complex. **(A)** Local resolution. **(B)** FSC curves. **(C)** Particle orientation distribution, and **(D)** Posterior directional distribution of particles.

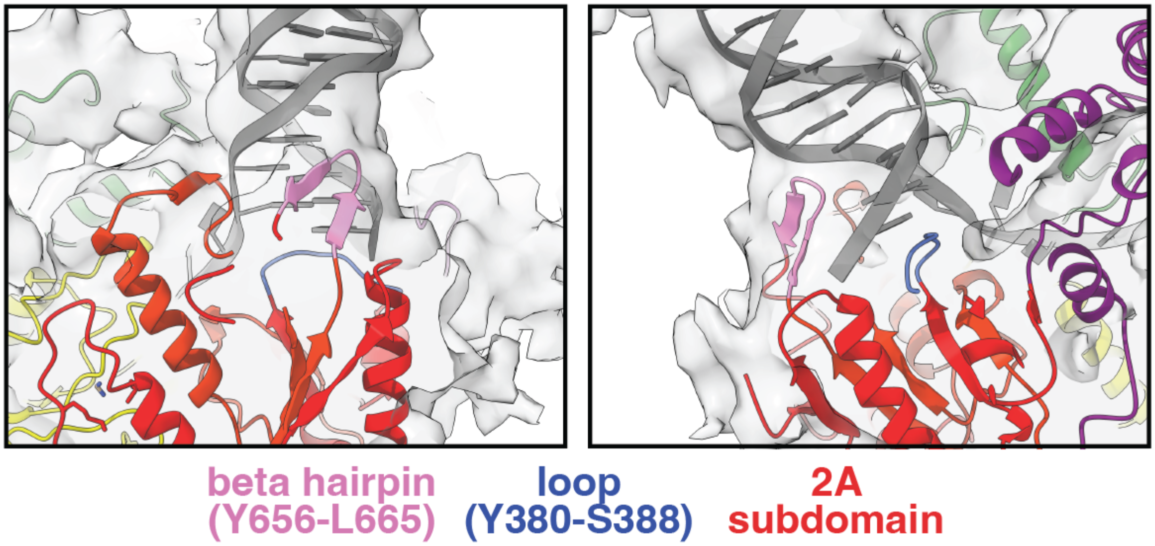


**Fig. S11:** Highlights of secondary structural elements of the 2A subdomain of the leading subunit contacting the DNA duplex and junction.

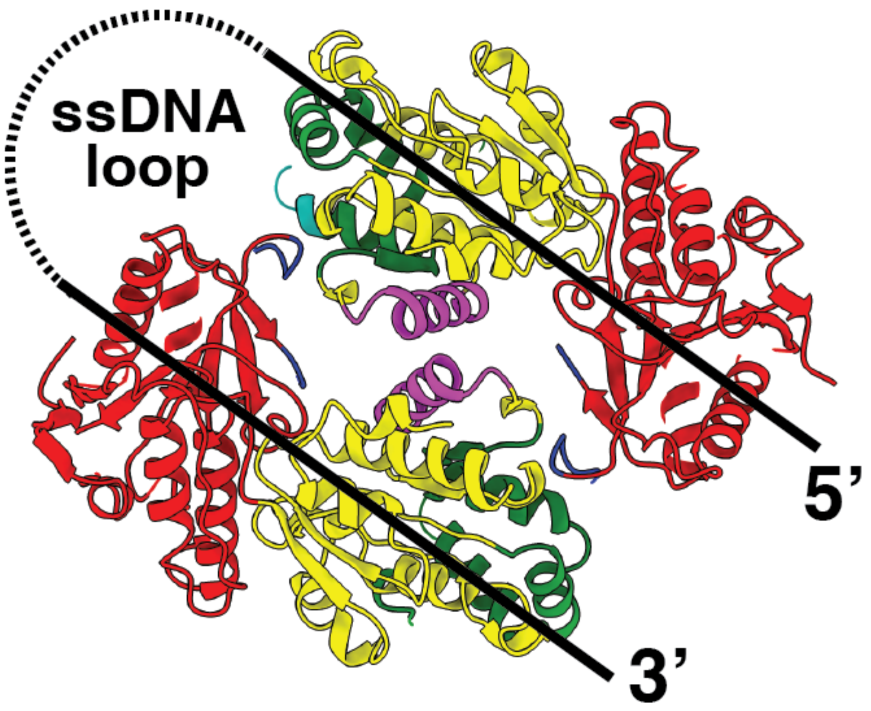


**Fig. S12:** A diagram indicating the need for a ssDNA loop if DNA were to bind to the apo compact motor conformation.

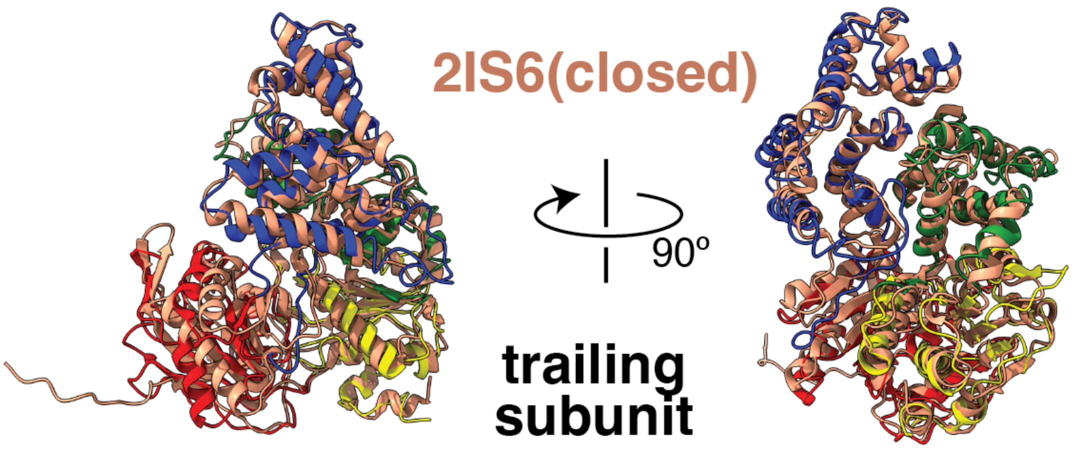


**Fig. S13:** Trailing subunit of *Mt*UvrD1 dimer-DNA complex aligned with closed *Ec*UvrD conformation (2IS6).

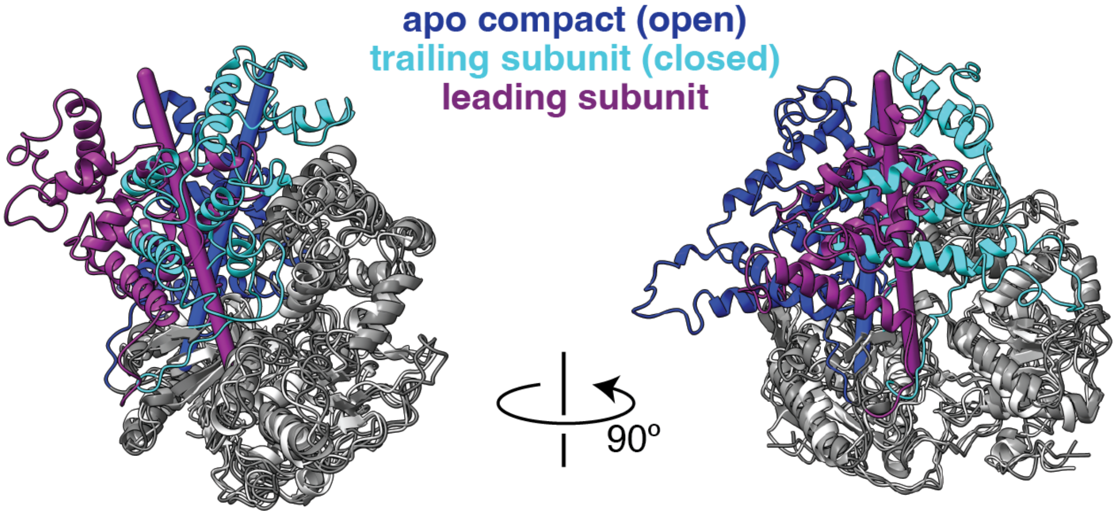


**Fig. S14:** Different axes of rotation describe the 2B movement from trailing to apo (*i.e.*, closed to open, blue to cyan, blue axis) and the 2B movement from trailing to leading (*i.e.*, closed to new conformation, purple to cyan, purple axis).

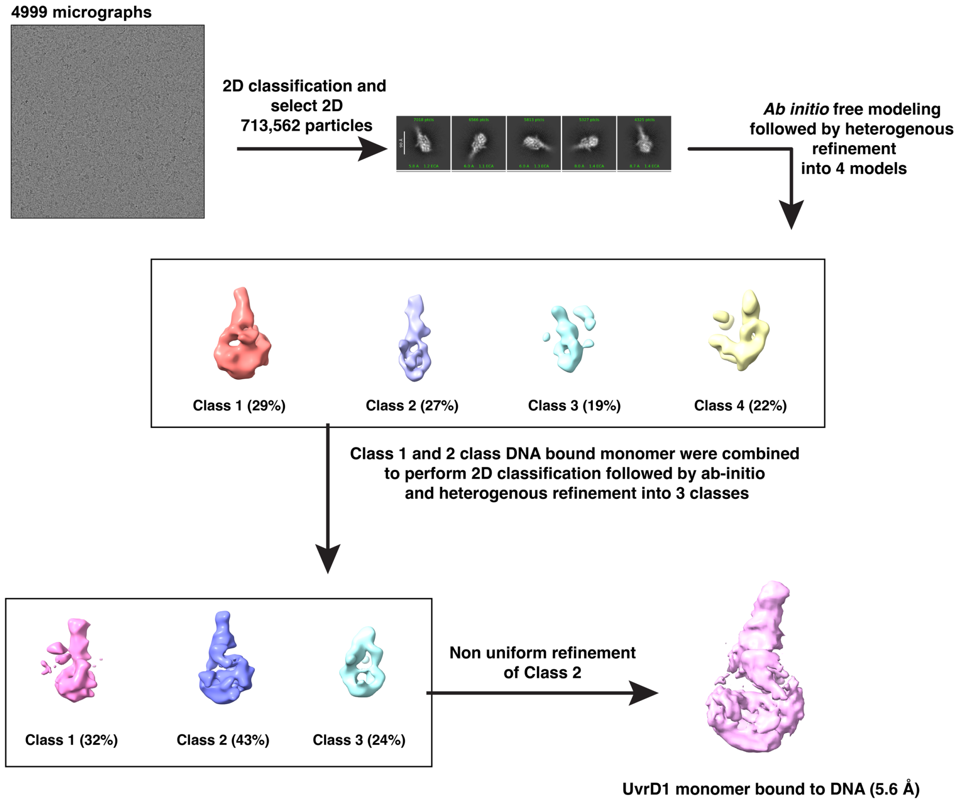


**Fig. S15:** Image processing pipeline for 5.9 Å DNA-bound *Mt*UvrD1 map.

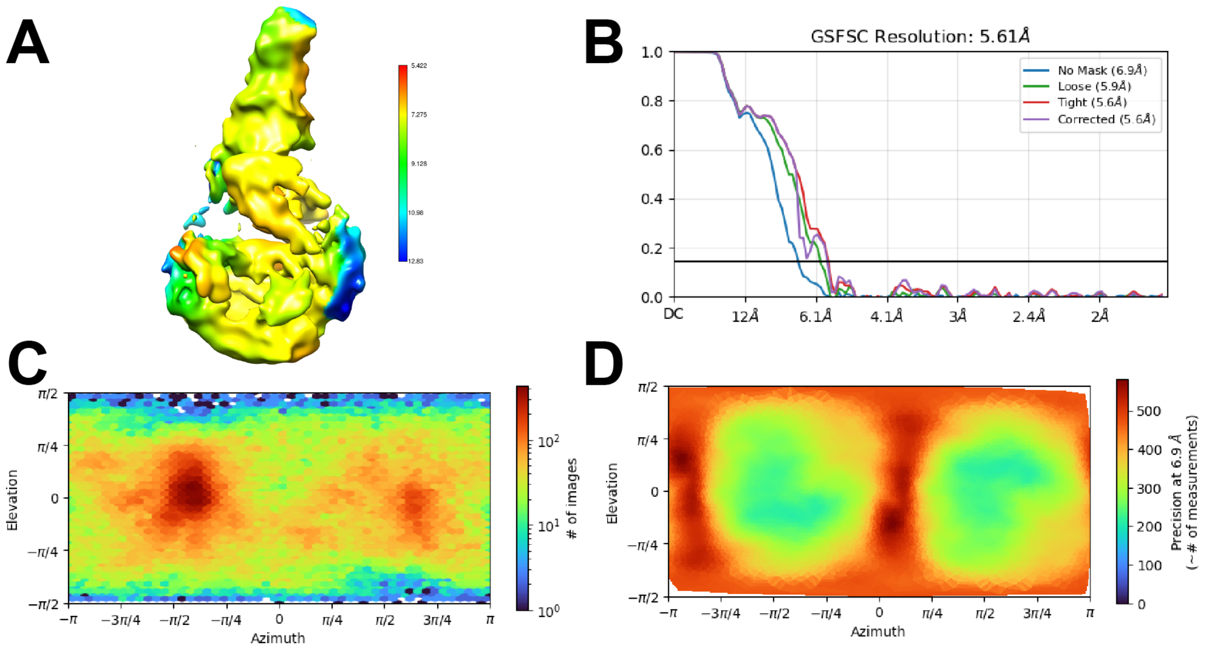


**Fig. S16:** Statistics for DNA-bound *Mt*UvrD1 monomer complex. **(A)** Local resolution. **(B)** FSC curve. **(C)** Particle orientation distribution, and **(D)** Posterior directional distribution of particles.

**Tables S1**

**Table S1:** Microscope parameters for apo dimer, DNA-bound dimer and DNA-bound monomer

| Parameter | Apo UvrD1 dimer | UvrD1 dimer with DNA | UvrD1 monomer with DNA |
|---|---|---|---|
| Microscope | Glacios | Glacios | Krios |
| Detector | Falcon IV and Gatan K3 | Falcon IV and Gatan K3 | CMOS and Gatan K2 |
| Magnification | 150,000 | 150,000 | 75,000 |
| Pixel size (A/pixel) | 0.94 Å/pixel | 0.928 Å | 0.868 Å |
| Total Dose | 46.46 e/Å$^2$ | 49.8 e/Å$^2$ | 59 e/Å$^2$ |
| Exposure time | 9.77 sec | 6.55 sec | 7.27 sec |
| Dose(electron/pix/sec) | 4.18e/pix/sec | 6.52e/pix/sec | 6 e/pix/sec |
| Fractions | 48 frames | 45 frames | 50 frames |
| Defocus range | 0.8 to -2.4 µ | -1 to -2.4 µ | -1 to -2.4 µ |
| Spherical aberration | 2.7 mm | 2.7 mm | 0.05 |
| Number of micrographs | 4562 | 4547 | 4999 |

**Movies S1 to S8:**

**Movie S1:** mS1_apocompact.mp4 shows the overall structure of the compact apo dimer of *Mt*UvrD1. Subdomains are colored as in the text and 2B-2B and 1A-1A dimer contacts are highlighted. See Fig. 2 of the main text.

**Movie S2:** mS2_DNAbound.mp4 shows the overall structure of the DNA-bound dimer of *Mt*UvrD1. The distinct 2B conformations are colored as in the text: Purple for the leading subunit and cyan for the trailing subunit. See Fig. 3 of the main text.

**Movie S3:** mS3_2ADNAcontacts.mp4 shows the details of the DNA contacts made by the 2A domain of the leading subunit of the DNA-bound dimer of *Mt*UvrD1. The beta-hairpin is in magenta, and the loop positioned where DNA unwinding appears to begin is in blue. See Fig. 3C of the main text.

**Movie S4:** mS4_2BrotationAxes.mp4 shows the distinct axes of rotation relating the 2B conformations of the apo compact (open, blue) and the leading dimer subunit (purple) to the trailing dimer subunit (closed, cyan). The axes are colored coded accordingly. See Fig. 3E of the main text.

**Movie S5:** mS5_ssDNAinterference.mp4 shows the clash between the path of the ssDNA through the leading dimer subunit and the 2B domain gating helix in the closed conformation (cyan). The gating helix in the leading dimer subunit is shown in purple. See Fig. 3F of the main text.

**Movie S6:** mS6_dsDNAinterference.mp4 shows the clash between the path of the dsDNA as held by the leading dimer subunit and the 2B domain in the open conformation (blue). See Fig. 3G of the main text.

**Movie S7:** mS7_MonomerComplex.mp4 shows the overall structure of a monomer of *Mt*UvrD1 bound to a DNA junction. See Fig. 4A of the main text.

**Movie S8:** mS8_LeadingSubunitMonomerComparison.mp4 shows the different relationship of the 2B domain to the DNA in the DNA-bound monomer (cyan) and the leading subunit of the DNA-bound dimer (purple, making contact with transparent purple 2B domain of the trailing subunit). See Fig. 4C of the main text.

**Movie S9:** mS9_ConformationMorph.mp4 visualizes the conformational change in the 2B subdomain between the DNA-bound monomer (cyan) and the DNA-bound dimer (purple). Residues contacting the DNA in the monomer structure are highlighted in yellow. See Fig. 4C of the main text.
